# Age-induced P-bodies become detrimental and shorten the lifespan of yeast

**DOI:** 10.1101/2021.11.05.467477

**Authors:** Joonhyuk Choi, Shuhao Wang, Yang Li, Nan Hao, Brian M. Zid

## Abstract

Aging is an irreversible process characterized by a progressive loss of homeostasis in cells, which often manifests as protein aggregates. Recently, it has been speculated that aggregates of RNA-binding proteins (RBPs) may go through pathological transitions during aging and drive the progression of age-associated neurodegenerative diseases. Using Saccharomyces cerevisiae as a model system of aging, we find that P-bodies —an RBP granule that is formed and can be beneficial for cell growth during stress conditions — naturally form during aging without any external stresses and an increase in P-body intensity is negatively correlated with the future lifespan of yeast cells. When mother cells transfer age-induced P-bodies to daughter cells, the mother cells extend lifespan, while the daughter cells grow poorly, suggesting that these age-induced P-bodies may be directly pathological. Furthermore, we find that suppressing acidification of the cytosol during aging slows down the increase in the intensity of P-body foci and extends lifespan. Our data suggest that acidification of the cytosol may facilitate the pathological transition of RBP granules during aging.

## Introduction

Aging can be defined as the progressive and irreversible decline in physiological function over time, arising from the loss of homeostatic mechanisms within the cell. One key function that declines in aging cells is protein homeostasis (proteostasis) as misfolded and aggregated proteins progressively accumulate. Protein aggregates are often manifested in many age-associated neurodegenerative diseases such as Alzheimer’s disease and amyotrophic lateral sclerosis (ALS) (1–3). One class of proteins that has been linked to a number of neurodegenerative diseases are RBPs. RBPs play important roles in regulating the interactions between RNAs and proteins, which can be mediated by intrinsically disordered regions contained in RBPs. As intrinsically disordered regions do not have a well-defined structure, RBPs are particularly sensitive to changes in physicochemical environment such as pH and temperature conferring a propensity for aggregation (4). When cells are young, they take advantage of the aggregation of RBPs to regulate gene expression during stress via formation of cytoplasmic messenger ribonucleoprotein (mRNP) granules. In stress conditions, cells localize mRNAs and proteins involved in RNA metabolism and translation into mRNP granules such as P-bodies and stress granules, presumably in order to protect cells from stresses and resume gene expression quickly when cells are free from stress conditions(5). Given that it was shown that P-bodies can be transferred to daughter cells in stress conditions and are beneficial for their growth(6), formation of mRNP granules can be beneficial to cells. Thus, during aging mRNP granules may initially form due to a decline in the ability to maintain homeostasis and yet confer some benefit for cellular survival at early ages. As cells get older, however, we hypothesize that these same mRNP granules may transition into a pathological state and become detrimental to cells.

Given the severe impacts of neurodegenerative diseases on quality of life, research has been done for several decades to find cures for them (7–9). However, we still do not understand the underlying causes for the occurrence of most neurodegenerative diseases, and thus there has been little progress toward curing them. In spite of the fact that aging is the major risk factor for neurodegenerative disease (10), as it is difficult to follow natural aging processes of humans, much of the research on neurodegenerative diseases has tried to mimic aging processes with genetic manipulations (11, 12) and has focused on understanding mutant forms of proteins, including RBPs, associated with the diseases (13–16). Mutations in the genes that encode these proteins are mainly linked with early-onset diseases, however, most cases of neurodegenerative diseases such as Alzheimer’s diseases, ALS, and Parkinson’s disease are late-onset and have no obvious genetic component(10). Therefore, it is thought that the prevailing cases of these diseases are associated with aging processes reflecting complex interactions between genetic and environmental factors. In order to address this perspective, we need to understand why and how wild type proteins transition to being pathological in an age-dependent manner.

Here, using *Saccharomyces cerevisiae* (budding yeast) as a model system of aging and tracking their natural aging processes over an entire lifespan in a microfluidics device (17), we observed that P-bodies naturally formed in yeast cells during aging without external stresses. Showing that these age-induced P-bodies go through a pathological transition over time and become detrimental to yeast cells, we provide evidence for our hypothesis that mRNP granules associated with the neurodegenerative diseases can become pathological to cells during aging. Our work suggests that an age-driven pathological transition of mRNP granules may contribute to the onset of age-related diseases.

## Results

### P-bodies naturally form during aging and become bigger and more intense over time

To measure the formation behaviors of mRNP granule components during aging in budding yeast, we used the Mother Enrichment Program (MEP) strain (18), a strain that was genetically modified to enrich mother cells in batch culture. We tagged the protein components of stress-induced mRNP granules such as P-bodies and stress granules with fluorescent proteins and measured the formation of fluorescent foci at different time points. We observed that the protein components of P-bodies such as Dcp2 (19) —the catalytic subunit of Dcp1-Dcp2 decapping enzyme complex— and Edc3 (20) —an enhancer of mRNA decapping— formed foci in 24 hours (Fig. 1A). Furthermore, these P-body protein components were mostly colocalized, suggesting that these age-induced foci were bona fide P-bodies (Fig. 1A and 1B). On the other hand, stress granule foci marked by Pab1-mNeonGreen were not observed during aging (Fig. 1A). As previously shown (21), Hsp104 foci —a disaggregase that is enriched in misfolded protein aggregates—also formed during early ages, but they were not colocalized with P-bodies (Fig. 1A). These observations suggest that age-induced formation of mRNP granules is specific to P-body components in budding yeast.

**Fig. 1.**
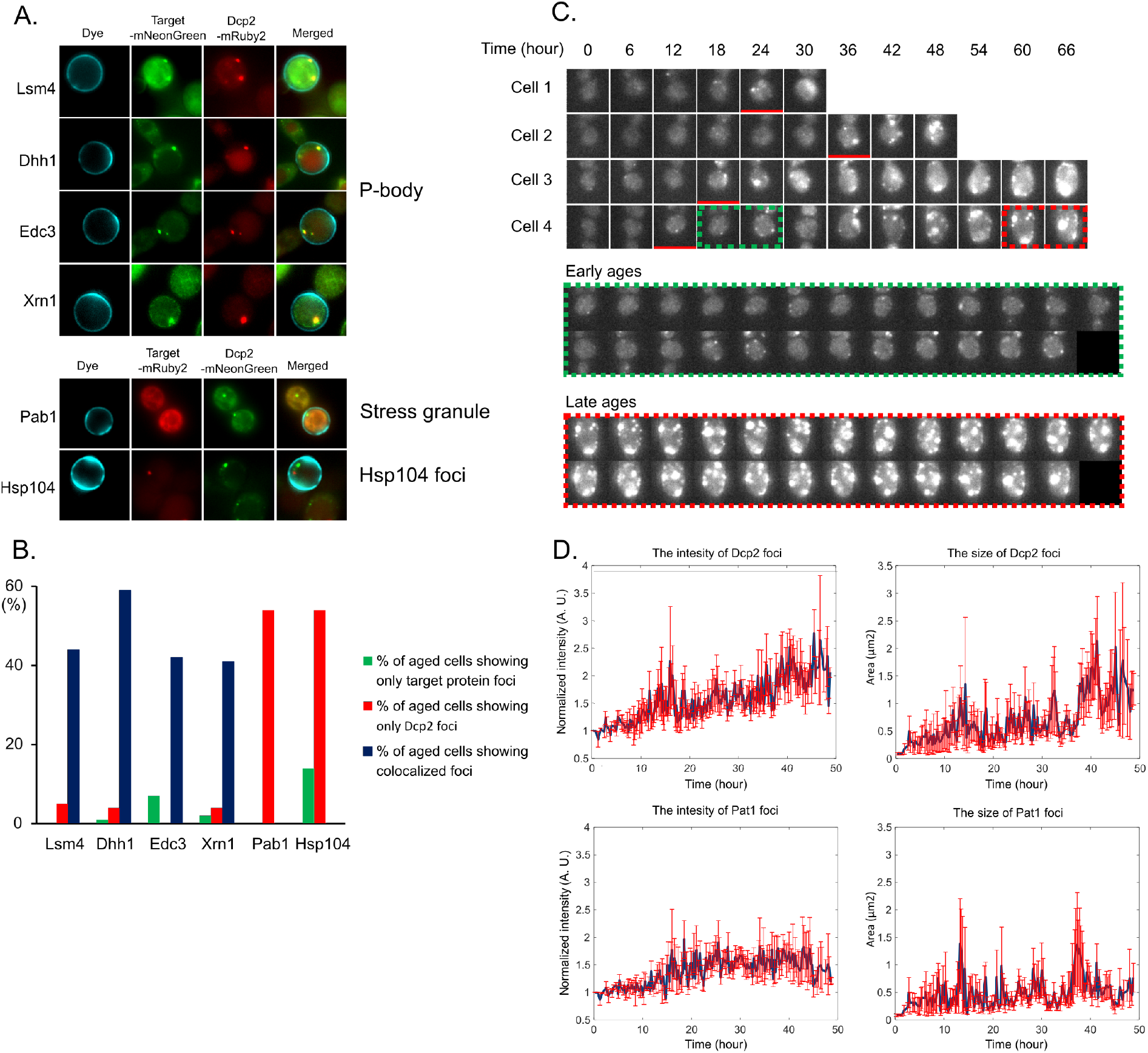
P-bodies naturally form during aging in the absence of external stress. A. Images of MEP strain cells that formed foci of P-body protein components or other mRNP granule markers (Pab1 and Hsp104). Cells were first stained with Dylight to indicate the mother cells before running aging experiments and images were taken 24 hours after starting the experiments. Note that Dcp2 foci were colocalized with only P-body protein components during aging. B. Quantification of colocalization distribution of Dcp2 foci and other protein components of mRNP granules. C. Representative images of wild type cells taken during their entire lifespans in a microfluidics device. Red underlines indicate when P-bodies were shown for the first time. The images in the red and green dashed boxes were obtained at every 15 minutes for 6 hours. Early aged cells indicated in green dashed boxes showed circular and transient foci. Late aged cells indicated in red dashed boxes showed highly intense and sustained foci. D. Changes in the intensities and the size of Dcp2-mRuby2 foci and Pat1-mNeonGreen foci taken from time course measurements in a microfluidics device (mean ± S. E., n=44). Error bars are indicated in red.

To track the age-induced formation of P-bodies across time, we tagged the P-body marker Dcp2 with mRuby2 in wild type cells and followed the formation of foci in a microfluidics device under normal growth conditions over their entire lifespans (Fig. 1C and 1D). As shown in the MEP strain, we observed that P-bodies formed during aging even in the absence of external stress. Following individual cells over time, we found that the first appearances of P-bodies among cells randomly occurred as cells aged. An interesting feature that we observed from the microfluidics device was that these P-bodies became larger and increase the intensities of their marker over time, which suggests that age-induced P-bodies are dynamic objects. When P-bodies appeared for the first time, they were small and were observed transiently over time. On the other hand, P-bodies found in late-aged cells were much larger and of higher intensity compared to those at their early ages, suggesting that the compositions of P-bodies can change once they form during aging (Fig. 1C).

In addition to Dcp2, we also measured the aggregation behaviors of another P-body protein Pat1 as well as proteins found in stress-induced aggregates such as Pab1 and Hsp104 in the microfluidics device (Fig. 1D and S1). We found that Pat1 (22) — a deadenylation-dependent mRNA-decapping factor enriched in P-bodies in stress conditions — also formed foci during aging and was colocalized with Dcp2 (Fig. S2). Interestingly, while the intensities and the sizes of Dcp2 foci kept increasing over time in general, the intensities and the sizes of Pat1 foci increased up to a certain level and then plateaued (Fig. 1D). As we found in the MEP strain (see Fig. 1A and 1B), we did not observe the formation of stress granules in wild type cells during aging (Fig. S1A). While Hsp104 foci formed at early ages, their sizes and intensities were not correlated with age over entire lifespans (Fig. S1B and S1C). These observations made in both the MEP strains and the wild type in the microfluidics device suggest that P-bodies naturally formed during aging even in the absence of external stresses while stress granules did not.

### An increase in protein components of age-induced P-bodies can act as a biomarker for future lifespan

As the intensities and the sizes of Dcp2 foci increased over time (Fig. 1D), we investigated whether the changes in Dcp2 foci are associated with aging phenotypes such as slow cell division rate during yeast lifespan. As the lifespan of each yeast cell is intrinsically heterogeneous, making it difficult to compare aging phenotypes among cells as a function of the number of cell division, we normalized the lifespan of each cell to their maximum number of cell divisions and compared its cell division rate with its normalized lifespan. We found that the division rates of mother cells were initially rather constant. After ~70% of their normalized lifespan, however, they decreased abruptly and rapidly (Fig. 2A). Furthermore, we found that the intensities of Dcp2 foci also increased rapidly at the normalized lifespan where cell division rate abruptly decreased ~70% of normalized lifespan). The intensity of Dcp2 foci at that point was 1.6 times higher than the threshold intensity for detecting Dcp2 foci (1.6x P-bodies). This relationship between cell division rate and the intensity of Dcp2 foci over time suggests that the changes in the intensity of Dcp2 foci can be associated with the occurrence of aging phenotypes.

**Fig. 2.**
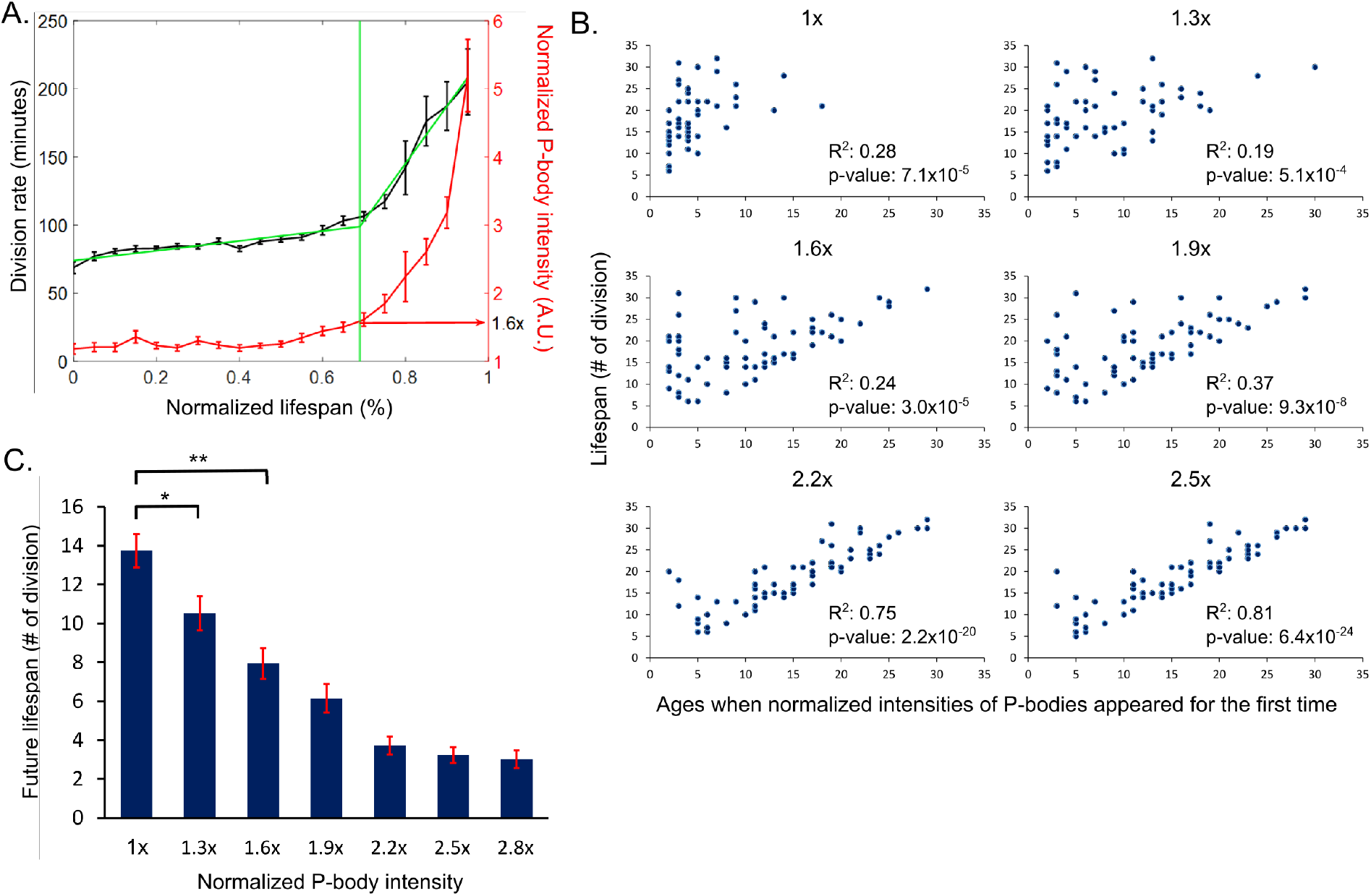
Formation of highly intense age-induced P-bodies is negatively correlated with future yeast lifespan. Formation of highly intense age-induced P-bodies is negatively correlated with future yeast lifespan. A. Division rate and normalized P-body intensity as a function of normalized lifespan (%). The green curve was obtained by fitting to a least squares spline curve with a degree of three. Note that the first appearance of P-bodies with an intensity of 1.6 times higher than the threshold intensity for detecting P-bodies (1.6x P-bodies) coincides with the time point at which the division rate increased abruptly. B. Plots of lifespan against the number of division at which each normalized intensity of P-bodies was shown for the first time. R2 and p-values for each panel were obtained by Spearman correlation fitting. Each dot corresponds to an individual cell. The intensities of P-bodies were normalized to the threshold at which P-bodies were detected in each cell. Note that the intensities of Dcp2 foci are positively correlated with the age of yeast. C. The plot of the future lifespan against the number of division at which each normalized intensity of P-bodies was shown for the first time (mean ± S. E.; two-sample t-test,*p=0.011, **p<10-5). Note that the intensities of Dcp2 foci are negatively correlated with the future lifespan of yeast.

To investigate how the changes in the intensity of Dcp2 foci are related to yeast lifespan, we compared the first occurrences of Dcp2 foci formation at various intensities to the lifespan of each cell (Fig. 2B). We found that timings at which P-bodies formed for the first time during aging was not significantly correlated with yeast lifespans. On the other hand, timings at which highly intense P-bodies formed for the first time during aging was significantly correlated with yeast lifespans; the higher the Dcp2-foci intensity observed, the better the correlations between the timing at which these foci formed for the first time and their lifespans were. In particular, when we measured the future lifespans of yeast cells after these age-induced P-bodies formed for the first time, the intensities of age-induced P-bodies were negatively correlated with the future lifespans of yeast cells (Fig. 2C). This behavior suggests that changes in the amount of protein components of the age-induced P-bodies can provide an insight into cellular ages such that Dcp2 intensities in P-bodies serve as a biomarker for future lifespan of yeast cells.

### Age-induced P-bodies become toxic to cells during aging

During aging we noticed that a small but significant portion of mothers with highly intense P-bodies transferred these granules to their daughter cells during replication as was previously seen during nutrient starvation conditions (6) (Fig. S3 and Supplementary Movie 1). Given that the intensities of P-body foci were negatively correlated with lifespan of yeast cells (Fig. 2B), we hypothesized that highly intense P-bodies could become toxic rather than beneficial during aging and an increase in the intensities of P-body foci could be an indication of this pathological transition. To investigate whether highly intense P-body foci formed during aging became toxic to cells, we measured changes in the division rate of daughter cells and the future lifespan of mother cells depending on the transfer status of highly intense P-bodies. If the highly intense P-bodies are toxic to cells, we expect to observe two things depending on their transfer status. First, given that daughter cells are rejuvenated relative to their mother cells unless their mother cells are very old (23), daughter cells that inherit the highly intense P-bodies from their mother cells would divide more slowly than those that do not if they were indeed toxic. Second, mother cells that remove the P-bodies by transferring them to daughter cells would live longer that those that do not. Given that the first appearance of 1.6x P-bodies coincides with the timing where yeast cells abruptly grow more slowly (Fig. 2A), we considered 1.6x the minimum intensity of P-bodies where they could be toxic. We subsequently refer to P-bodies whose normalized intensities are higher and lower than 1.6x as “intense” and “weak” P-bodies, respectively, and investigated the effects of transfer of these intense P-bodies on the division rate of daughter cells and the future lifespan of mother cells.

First, we found that the division rates of daughter cells slowed down only when the intense P-bodies were transferred to them (Fig. 3A and 3B). When daughter cells inherited the intense P-bodies from their mother cells, their division rates ~418 minutes were much slower than normal young cells (90 minutes). Furthermore, no further cell divisions or very long division rates (more than 6 hours) were observed in more than 80% of the daughter cells (Fig. 3B). When daughter cells did not receive the intense P-bodies from their mother cells, on the other hand, their division rates ~92 minutes) were similar to the division rate of normal young cells. These observations suggest that the transferred intense P-bodies were toxic enough to interfere significantly with the cell division of daughter cells. In the case of the weak P-bodies, the division rates of daughter cells that inherited those P-bodies ~115 minutes) were not significantly slower than those that did not ~99 minutes, Fig. 3C), suggesting that the weak P-bodies appear less toxic to cells than the intense P-bodies. To rule out a possibility that these differences in the division rates of daughter cells result from different ages of mother cells for the transfer and non-transfer cases, we compared the average ages of mother cells in analyzing the transfer and non-transfer cases and found that there was no difference in the average ages of mother cells between them (Table S2). Therefore, it is unlikely that these different division rates of daughter cells that depend on the transfer status of P-bodies result from different ages of mother cells.

**Fig. 3.**
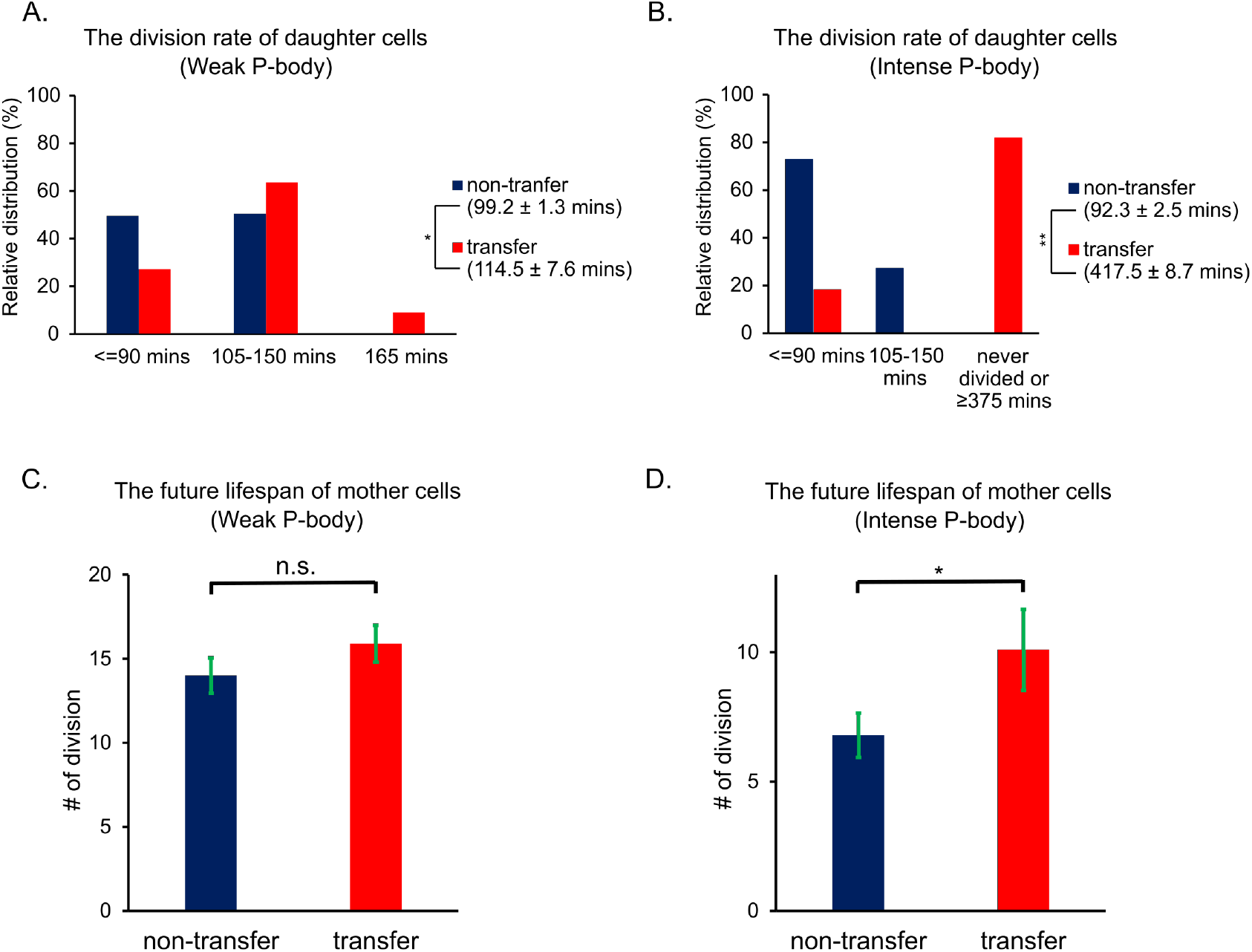
Age-induced P-bodies go through pathological transitions during aging and become detrimental to cells. A-B. The division rates of daughter cells depending on whether mother cells transferred the weak or the intense P-bodies to their daughter cells, respectively (mean ± S. E.; two-sample t-test for the weak P-body cases, *p=0.0015, two-sample t-test for the intense P-body cases, **p<10-10; number of cells, non-transferred (n=113) and transferred (n=11) for the weak P-bodies, non-transferred (n=59) and transferred (n=11) for the intense P-bodies). The daughter cells that were dead before dividing for the intense P-body cases were excluded when growth rates were averaged. Note that more than 80% of daughter cells that inherited the intense P-bodies from their mother cells never divided in the time of viewing (at least 6 hours and longer) or were dead before dividing, while all daughter cells that inherited the weak P-bodies went through cell divisions. C-D. The future lifespans of mother cells depending on whether mother cells transferred the weak or the intense P-bodies to their daughter cells (mean ± S. E.; two-sample t-test, not significant (n.s.), *p<0.05; number of cells, non-transferred (n=38) and transferred (n=12) for the weak P-bodies, non-transferred (n=43) and transferred (n=24) for the intense P-bodies).

In addition, we also found that the future lifespan of mother cells was extended when the intense P-bodies were removed from them (Fig. 3C and 3D). When mother cells transferred the intense P-bodies to their daughter cells, they lived longer than those that did not (Fig. 3D and S3). On the other hand, transferring the weak P-bodies to daughter cells did not significantly affect the future lifespan of mother cells (Fig. 3C). While P-bodies formed in nutrient starvation conditions are beneficial for cell growth (6), these data suggest that age-induced P-bodies may go through pathological transitions over time and the intense P-bodies may drive pathology in aged cells.

### Overexpression of an assembly unit of vacuolar ATPase represses cytosolic acidification during aging and suppresses the pathological transition of P-bodies

It was previously shown that cytosolic pH decreases with aging in yeast (24, 25). As phase separation of proteins is pH-sensitive (26–29) and specifically intrinsically disordered regions in Dcp2 are more sensitive to pH perturbation than well-defined domains (30, 31), we hypothesized that a decrease in cytosolic pH during aging could contribute to the rise in formation of P-bodies and even facilitate the pathological transition of P-bodies. It was shown that vacuolar pH increases during aging as opposed to cytosolic pH and the increase in vacuolar pH can be suppressed by overexpressing Vph2, a protein required for assembly of vacuolar H+-ATPases (32, 33). Given that the cytoplasm is a proton source for vacuolar H+-ATPases, we tested whether overexpression of *VPH2* would also delay acidification of the cytosol during aging. To test this hypothesis, we measured changes in cytosolic pH levels during aging with pHluorin in wild type cells and in a strain that overexpresses *VPH2* (*VPH2* OE) using a microfluidics device over time. Ratiometric pHluorin is a pH-sensitive GFP variant that displays a bimodal excitation spectrum with peaks at 395 and 475 nm and an emission maximum at 509 nm (34). Upon acidification, pHluorin excitation at 395 nm decreases with a corresponding increase in the excitation at 475 nm such that the distinctive 395/475-nm excitation ratio (pHluorin ratio) decreases. We found that initially there was no difference in cytosolic pH levels between wild type cells and *VPH2* OE cells and cytosolic pH levels in both strains decreased to the similar levels when we measured them at 24 hours (Fig. 4A and 4B). After 48 hours, on the other hand, cytosolic pH levels in wild type cells kept decreasing, while cytosolic pH levels in *VPH2* OE cells did not decrease to the same extent and instead were similar to the pH levels measured at 24 hours (Fig. 4A and 4B). Therefore, we conclude that overexpression of *VPH2* suppresses acidification of the cytosol during the later stages of aging.

**Fig. 4.**
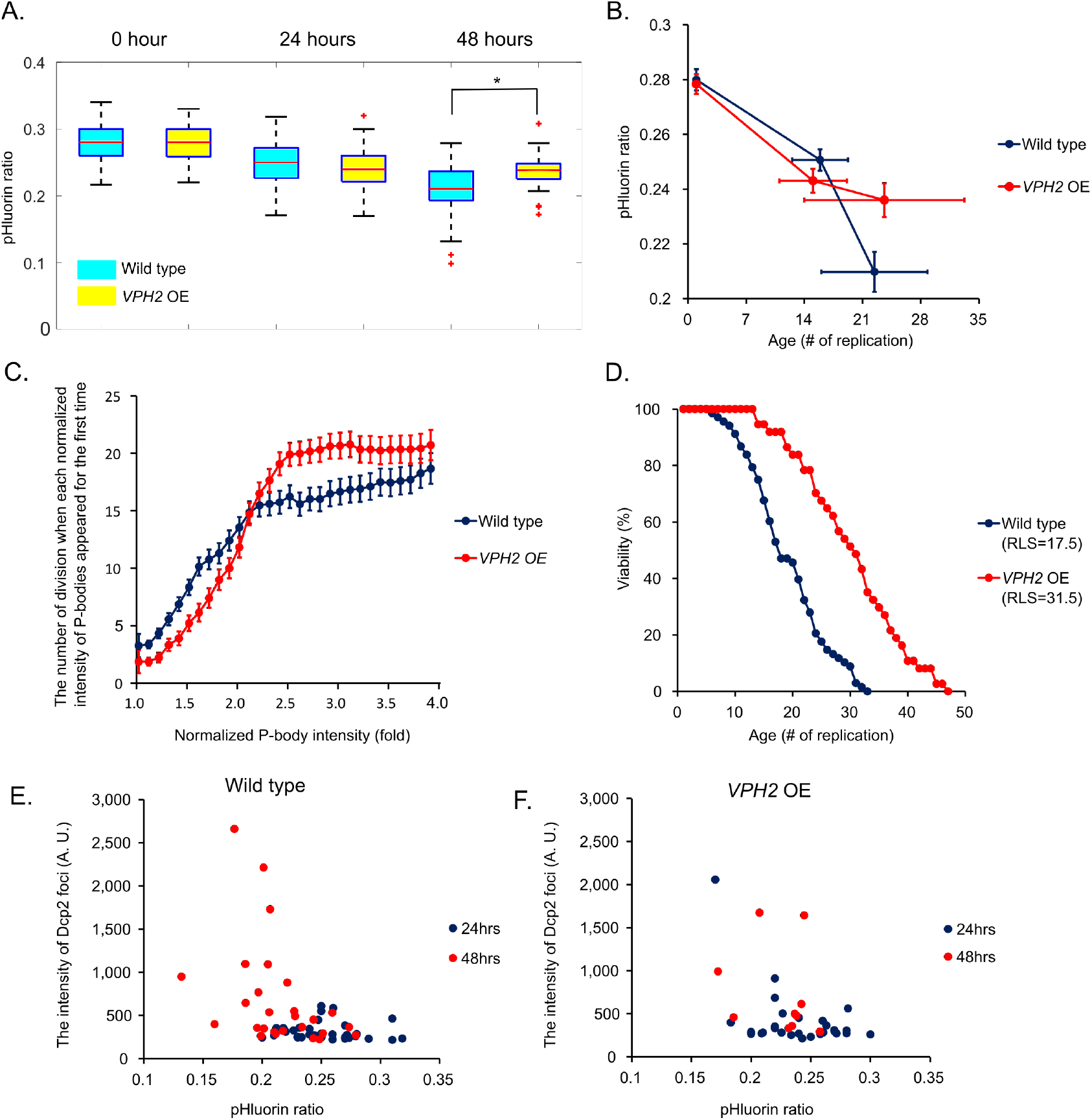
Overexpression of an assembly unit of vacuolar ATPase represses cytosolic acidification during aging and suppresses the pathological transition of P-bodies. A. The box plots of pHluorin ratiometric values in the wild type and *VPH2* OE at 0, 24 and 48 hours after aging experiments were initiated in a microfluidics device (two-sample t-test, *p=0.012). pHluorin emission intensities were obtained by dual excitation at 405 nm and 485 nm with an emission filter of 535 nm. Ratios of 405 nm/485 nm were shown as pHluroin ratios. On each boxplot, the central red line indicates the median and the bottom and tope edges of the box indicate the 25th and the 75th percentiles, respectively. Red “+” symbols indicate the outliers. B. Averaged pHluorin ratios obtained from Figure 4A at each time. Cellular ages were converted into the number of divisions. C. The number of division when each normalized intensity of P-bodies appeared for the first time as a function their intensities normalized to the detection threshold of individual cells. D. Lifespan analysis of wild type cells and *VPH2* OE cells. E-F. The plot of the intensity of Dcp2 foci vs pHluorin ratio at 24 and 48 hours in wild type cells and *VPH2* OE cells, respectively.

We next tested whether the suppression of cytosolic acidification via *VPH2* overexpression would also delay the formation of highly intense Dcp2 foci. To test this hypothesis, we compared the first timing of formation of Dcp2 foci with different intensities between wild type and *VPH2* OE cells. We found that initially Dcp2 foci formed in VPH2 OE cells earlier than in wild type cells (the 2nd and the 3rd division, respectively) and intensities of Dcp2 foci in *VPH2* OE cells were higher than those in wild type cells up to ~15 generations (Fig. 4C). We noted that until ~15 generations cytosolic pH levels in *VPH2* OE cells were slightly lower than pH levels in wild type cells (Fig. 4B). After ~15 generations, however, the intensities of Dcp2 foci in *VPH2* OE cells increased more slowly during aging than those in wild type cells. These trends in wild type cells and *VPH2* OE cells during aging resemble the changes in pH during aging (Fig. 4B). We also observed that *VPH2* OE cells lived ~50% longer than wild type cells as previously reported (32) (Fig. 4D), which is consistent with our hypothesis that delaying the formation of highly intense P-bodies would extend lifespan. These observations suggest that slow acidification in the *VPH2* OE strain may suppress the increase in the intensity of Dcp2 foci during aging.

Furthermore, we found that the intensities of Dcp2 foci were negatively correlated with cytosolic pH levels (Fig. 4E and 4F). At 24 hours where pH levels in wild type and *VPH2* OE cells were similar to each other, pHluorin ratios in both strains mostly lay between 0.2 and 0.3 and the intensities of Dcp2 foci in these cells were similar to each other regardless of their pHluorin ratios. At 48 hours where pH levels in wild type cells were lower than those in *VPH2* OE cells, on the other hand, ~40% of wild type cells exhibited pHluorin ratios below 0.2 and the intensities of Dcp2 foci in these cells were much higher than those whose pHluorin ratios were above 0.2. Most cells of *VPH2* OE still exhibited pHluorin ratios above 0.2 such that the differences in the intensities of Dcp2 foci in *VPH2* OE between 24 and 48 hours were much smaller than the differences in wild type cells. Therefore, these observations suggest that cytosolic acidification during aging can contribute to the formation and pathological transition of P-bodies.

## Discussion

The majority of cases of neurodegenerative diseases are associated with general aging processes rather than specific genetic mutations in disease-associated proteins. In order to understand the underlying causes for the occurrence of most neurodegenerative diseases, it is important to understand how wild type proteins transition to being pathological over the natural aging process. Using a microfluidics device that allows us to measure natural aging processes of yeast cells over their entire lifespans, we found that P-bodies naturally formed during aging and they underwent a pathological transition making them toxic to cells over time. While many agerelated changes have been cataloged across organisms, for most of the changes it is difficult to ascribe whether they are directly pathological and driving aging, or just a byproduct of aging (35). Because highly intense P-bodies sometimes naturally transferred from mother to daughter cells, we were able to observe the impact of (i) loss of highly intense Pbodies on the lifespan of mother cells and (ii) gain of highly intense P-bodies on the division rate of daughter cells and could therefore determine that highly intense P-bodies are directly pathological. Given that the pathological transition of P-bodies naturally happened during aging, our finding implies that the transition of mRNP granules may drive the pathology of aging rather than just being a byproduct from age-associated processes.

The pathological transition of mRNP granules during aging may be a conserved phenomenon in higher organisms. It was shown that P-bodies form during aging in many organisms including yeast cells and *C. elegans* (21, 36, 37). For example, Rieckher et al. showed in C. elegans that the concentration of the protein components of P-bodies increased during aging and that the translation initiation factor eIF4E isoform IFE-2 was increasingly sequestered in P-bodies during aging(37). In addition, Lechler, et al. showed that key stress granule (SG) proteins PAB-1 and TIAR-2 form aggregates in aged C. elegans and a high level of aggregation of these SG components is associated with shorter lifespan (35). Given that P-bodies and SGs are conserved from yeast to humans, it is plausible that the transition of mRNP granules may drive the pathology of aging in higher organisms.

Furthermore, our finding that the transition of P-bodies may drive the pathology of aging process highlights P-bodies as a promising target of research on neurodegenerative diseases. Because many proteins associated with neurodegenerative diseases such as TDP-43 and FUS are localized to stress granules (SGs) in patients, much of the research has focused on SGs as a cause of disease pathology. However, there has been growing evidence that these proteins are also localized in P-bodies. For example, it was shown that TDP-43 was colocalized with P-bodies in ALS motor neurons, but not in normal neurons, leading to sequestration of low molecular neurofilament mRNA to P-bodies in ALS neurons (38). When FUS was expressed in yeast, it induced the formation of P-bodies as well as SGs and was colocalized with P-bodies(39). As docking and fusion of P-bodies with SGs can happen in mammalian cells (40, 41), the transition of P-bodies that sequester the disease-associated proteins during aging may contribute to the pathology of the age-associated neurodegenerative diseases along with mislocalization of the proteins to SGs.

We show that suppressing cytosolic acidification during aging can lead to delaying the pathological transition of age-induced P-bodies (Fig. 4). These findings are of particular interest because a decrease in cytosolic pH during aging was also observed in other higher organisms such as *D. melanogaster* and human cells (42, 43). Furthermore, from the perspective of chronological lifespan defined as the length of time that a non-dividing yeast cell survives in the stationary phase (44), acidification of the extracellular environment limits the chronological lifespan of yeast cells and survival of quiescent human cells in culture4 (45, 46). Therefore, it is plausible that acidification of the cellular environment during aging could be a general driving force behind the formation of mRNP granules and facilitates their pathological transition in humans. In this regard, finding a way to suppress abnormal acidification of cytoplasm during aging could be useful to develop possible remedies for neurodegenerative diseases.

Mechanistically, it remains unclear how these highly intense P-bodies become toxic to cells during aging. As the protein concentration enriched in P-bodies is inversely correlated with cytoplasmic exchange rate (47), highly intense P-bodies may lead to dysregulation of gene expression due to a large difference in the exchange rate of mRNAs and proteins enriched in the P-bodies with the cytoplasm. Furthermore, we observed that the relative compositions of Pat1 and Dcp2 enriched in P-bodies changed over time (Fig. 1D). Therefore, it is plausible that RNA metabolism occurring within P-bodies may deteriorate during aging due to larger differences in the stoichiometry between Dcp2 and Pat1 or perhaps other protein components in P-bodies.

In summary, our work suggests that mRNP granules that naturally form during aging may go through pathological transitions and drive the progression of age-associated diseases. Further determination of molecular mechanisms by which mRNP granules make a pathological transition will provide an insight into age-associated neurodegenerative diseases and could inform the development of novel therapeutic approaches to treat them.

## ACKNOWLEDGEMENTS

We thank the Zid lab for feedback on this research and manuscript, especially Anna Guzikowski. We would like to thank Catherine Triandafillou from the Drummond lab for providing the pHluorin plasmid and the Denic Lab for providing the MEP strain. This work was supported by an AFAR Junior Faculty Grant (to BMZ), National Institutes of Health R21AG064342 (to BMZ), Hellman Fellows Fund (to BMZ), R01GM111458 (to NH), R01AG068112 (to NH) and R01AG056440 (to NH).

## Author Contributions

J. C., S. W., and Y. L. performed the experiments; J.C. analyzed the data; J.C. and B. M. Z wrote the original manuscript; N. H. reviewed and edited the original manuscript

## Competing interests

The authors declare no competing financial interests.

## Methods

### Strain construction

All strains used in this study were in the BY4741 (*MATa his3*Δ*1 leu2*Δ*0 met15*Δ*0 ura3*Δ*0*) strain background and they were listed in Supplementary Table 1. A strain harboring NHP6a-iRFP-kanMX was a gift from the Hao lab at UCSD. Plasmids used to make these strains were listed in Supplementary Table 1. pCGT05-pTDH3-pHluorin-LEU plasmid used to express pHluorin in yeast strains was a gift from the Drummond lab at the University of Chicago. Transformations were performed with a high-efficiency version of the lithium acetate/single-stranded carrier DNA/PEG method48. The colonies of transformation were confirmed by PCR and/or phenotype screening such as measuring fluorescence signals from colonies.

### Aging experiments with MEP strains

Cells were grown overnight in synthetic complete (SC) media with 2% glucose at 30°C. The overnight culture was inoculated and was grown for at least 16 hours at 30°C. During cell growth, the final OD600 was always below 0.3. To label the cell wall of mother cells, they were harvested by centrifugation at 3000g of rcf for 5 minutes and were washed two times with 1 ml of pre-warmed 1x PBS buffer with 2%glucose at 30°C. Cell pellet was resuspended with 1ml of 1μg/ml of Dylight633 dye in 1x PBS buffer with 2% glucose and was incubated for ~15 minutes at 30°C. The labeled mother cells were washed twice with pre-warmed SC media with 2%glucose at 30°C and were inoculated to fresh SC media with 2% glucose such that the final OD600 = 0.003. After the inoculated cell culture recovered for 1 hour at 30°C under shaking (220rpm), β-estradiol (Sigma-Aldrich) was added to the recovered cell culture such that the final concentration is 1μM. In 24 hours, the cell culture was spun down at 3000g of rcf and was resuspended in 20 μl. 2μl of resuspended cell culture was put on the glass slide to take images. Images taken from Cy5 channel were used to identify labeled old mother cells and colocalization analysis was performed with images taken from GFP and RFP channels.

### Cell growth and media for microfluidics device experiments

Cell growth conditions for microfluidics device experiments were the same as the ones published from the Hao lab at UCSD (17, 48, 49). Before cells were loaded into the microfluidics device, they were grown in SC media with 2% glucose at 30 °C up to OD600 = 0.2 at least for 12 hours. Then, the overnight culture was inoculated such that the final cell culture was grown for at least 16 hours up to OD600 = 0.2. After cells were loaded into the device, they were grown in SC media with 2% glucose and 0.04% Tween-20 at 30°C.

### Time-lapse microscopy in a microfluidics device

All time-lapse microscopy was performed on a Nikon Ti-E microscope with a 60× Plan Apo Lambda objective and Perfect Focus System (PFS, Nikon). Fluorescence was excited by an LED light source (Sola, Lumencor) and collected to an sCMOS camera (PCO Edge 4.2Q). The excitation and emission filters for GFP, RFP and pHluorin are: Ex 480/30nm Em 460/50, Ex 560/40nm Em 635/60nm, and Ex 395/25nm Em 510 nm long-pass, respectively. Automated microscopy was conducted with NIS-Elements and the temperature was maintained at 30°C with a customized heating chamber. The design of microfluidics device and the protocol for setting up the microfluidics experiments were obtained from Li et al (17). Our microfluidics devices were modified from the original design such that the width of chamber for loading cells can be either 6.5μm or 6μm in order to harbor cells bigger than wild type cells.

### Cell segmentation and analysis of Dcp2-mRuby2 foci

Image analysis was performed with custom Matlab (Mathwork) codes. For cell segmentation, at first cells located in the dent of the trap in the chamber were labeled. The labeled cells were segmented from bright-field images using watershed transformation at each time frame. To track the labeled cells over time, the center of mass of each segmented region was calculated at evert time frame and each labeled cell at the next time frame was identified as the one with a minimal distance from the center of mass of the labeled cell at the previous time frame. To identify P-bodies indicated by Dcp2-mRuby2 within the labeled cells, first background-subtracted RFP images were segmented using the mask obtained from the bright-field images and Dcp2 foci within the segmented region were identified by finding pixels whose values exceeded an empirically defined threshold. The intensities of P-bodies were calculated by the mean of top 9 pixel values within identified Dcp2 foci. Cell divisions of each mother and daughter cell were manually identified and were counted when mother and daughter cells were physically separated in bright-field images.

### Determination of the status of P-body transfer from a mother cell to a daughter cell

To determine whether P-bodies in a mother cell were transferred to its daughter cell, the maximum intensity of P-bodies in mother and daughter cells were compared after the intensities of P-bodies in each cell over their entire lifespan were obtained. The same threshold intensities that we used in their mother cells were applied in order to identify P-bodies in daughter cells. To decide if P-bodies were transferred from a mother cell to a daughter cell at the n-th frame, following criteria should be satisfied; 1) there was no P-body identified in a daughter cell at the (n-1)th and the n-th frame, 2) the maximum intensity of P-bodies found in the mother cell at the (n+1)th frame was smaller than 0.6 times of the maximum intensity of P-bodies in the mother cell at the n-th frame, and 3) the maximum intensity of P-bodies found in the daughter cell at the (n+1)th frame was higher than 0.7 times of the maximum intensity of P-bodies in the mother cell at the n-th frame. If no P-bodies were identified at the (n+1)th frame in a mother cell, which is most of the detected cases in Figure 3, only condition 1) and 3) above were applied.

## Notes

### Competing Interest Statement

The authors have declared no competing interest.

