## Supplementary material for "Age-induced P-bodies become detrimental and shorten the lifespan of yeast": Figure Supplements + Tables

| Strain Name | Description |
|---|---|
| ZY101 | *MEP strain, BY4741 MATα ade2::hisG his3 leu2 trp1Δ63 ura3Δ0 met15Δ::ADE2 hoΔ::PSCW11-cre-EBD78-NATMX loxP-UBC9-loxP-LEU2 loxP-CDC20-Intron-loxP-HPHMX, Dcp2-mNeonGreen-HIS, Pab1-mRuby2-URA* |
| ZY103 | *MEP strain, BY4741 MATα ade2::hisG his3 leu2 trp1Δ63 ura3Δ0 met15Δ::ADE2 hoΔ::PSCW11-cre-EBD78-NATMX loxP-UBC9-loxP-LEU2 loxP-CDC20-Intron-loxP-HPHMX, Dcp2-mNeonGreen-HIS, Hsp104-mRuby2-URA* |
| ZY117 | *MEP strain, BY4741 MATα ade2::hisG his3 leu2 trp1Δ63 ura3Δ0 met15Δ::ADE2 hoΔ::PSCW11-cre-EBD78-NATMX loxP-UBC9-loxP-LEU2 loxP-CDC20-Intron-loxP-HPHMX, Dcp2-mRuby2-URA, Lsm4-mNeonGreen-HIS* |
| ZY133 | *MEP strain, BY4741 MATα ade2::hisG his3 leu2 trp1Δ63 ura3Δ0 met15Δ::ADE2 hoΔ::PSCW11-cre-EBD78-NATMX loxP-UBC9-loxP-LEU2 loxP-CDC20-Intron-loxP-HPHMX, Dcp2-mRuby2-URA, Dhh1-mNeonGreen-HIS* |
| ZY134 | *MEP strain, BY4741 MATα ade2::hisG his3 leu2 trp1Δ63 ura3Δ0 met15Δ::ADE2 hoΔ::PSCW11-cre-EBD78-NATMX loxP-UBC9-loxP-LEU2 loxP-CDC20-Intron-loxP-HPHMX, Dcp2-mRuby2-URA, Edc3-mNeonGreen-HIS* |
| ZY135 | *MEP strain, BY4741 MATα ade2::hisG his3 leu2 trp1Δ63 ura3Δ0 met15Δ::ADE2 hoΔ::PSCW11-cre-EBD78-NATMX loxP-UBC9-loxP-LEU2 loxP-CDC20-Intron-loxP-HPHMX, Dcp2-mRuby2-URA, Xrn1-mNeonGreen-HIS* |
| ZY250 | *BY4741 MATa his3Δ1 leu2Δ0 ura3Δ0 met15Δ0, NHP6a-iRFP-kanMX, Dcp2-mRuby2-HPH, Pat1-mNeonGreen-URA* |
| ZY272 | *BY4741 MATa his3Δ1 leu2Δ0 ura3Δ0 met15Δ0, Dcp2-mRuby2-NAT, Pab1-mNeonGreen-HIS* |
| ZY652 | *BY4741 MATa his3Δ1 leu2Δ0 ura3Δ0 met15Δ0, NHP6a-iRFP-kanMX, Dcp2-mRuby2-HIS, Hsp104-mNeonGreen-URA* |
| ZY653 | *BY4741 MATa his3Δ1 leu2Δ0 met15Δ0, NHP6a-iRFP-kanMX, Dcp2-mRuby2-HIS, ura3Δ::PGPD-VPH2-tCYC1-URA* |
| ZY654 | *BY4741 MATa his3Δ1 met15Δ0, NHP6a-iRFP-kanMX, Dcp2-mRuby2-HIS, leu2Δ::PTDH3-pHluorin-LEU,ura3Δ::PGPD-VPH2-tCYC1-URA* |
| ZY655 | *BY4741 MATa his3Δ1 leu2Δ0 ura3Δ0 met15Δ0, NHP6a-iRFP-kanMX, Dcp2-mRuby2-HIS, leu2Δ::PTDH3-pHluorin-LEU* |

| Plasmid Name | Description |
|---|---|
| ZP46 | pKT-mNeonGreen-HIS |
| ZP217 | pKT-mRuby2-HIS |
| ZP109 | pKT-mRuby2-HPH |
| ZP112 | pKT-mRuby2-NAT |
| ZP47 | pKT-mNeonGreen-URA |
| ZP309 | pAG306-PGPD-VPH2-tCYC1 chrI |
| ZP485 | pCGT05-PTDH3-pHluorin-LEU |

**Intense P-body**

| | observation frequency (%) (average number of divisions of mother cells) | |
|---|---|---|
| | Transfer (n=11) | Non-transfer (n=59) |
| <=90 mins | 18.18% (12) | 72.9% (15.1) |
| 105-150 mins | | 27.1% (18.3) |
| >=375 mins | 36.36% (12.5) | |
| never divided | 45.45% (15.2) | |
| total averaged age of mother cells | 13.6 | 15.9 |

**Weak P-body**

| | observation frequency (%) (average number of divisions of mother cells) | |
|---|---|---|
| | Transfer (n=11) | Non-transfer (n=113) |
| <=90 mins | 27.27% (11) | 49.56% (11) |
| 105-150 mins | 63.64% (12.9) | 50.44% (13.8) |
| 165 mins | 9.09% (12) | |
| total averaged age of mother cells | 12.3 | 12.4 |

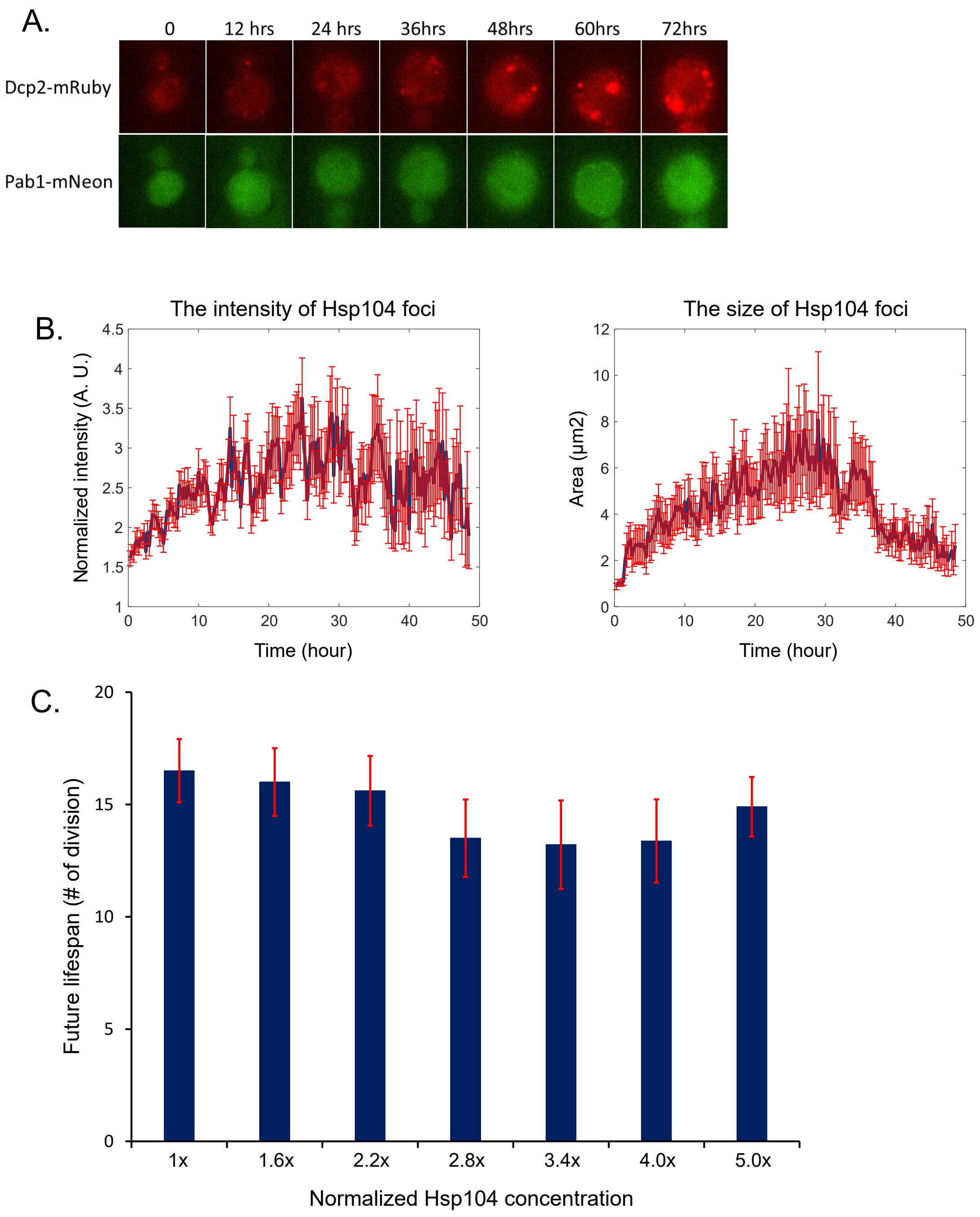


**Figure S1. Formation of protein foci other than P-bodies in wild type cells in a microfluidics device during aging. A.** Time course images of wild type cells harboring Dcp2-mRuby2 and Pab1-mNeonGreen. Note that no Pab1 foci were shown over the time course. **B.** Changes in the intensities and the foci size of Hsp104-mNeonGreen over time in wild type cells during aging in a microfluidics device (mean ± S. E.). Red bars indicate error bars. **C.** The plot of the future lifespan and the number of division at which each normalized intensity of Hsp104 foci was shown for the first time (mean ± S. E.).

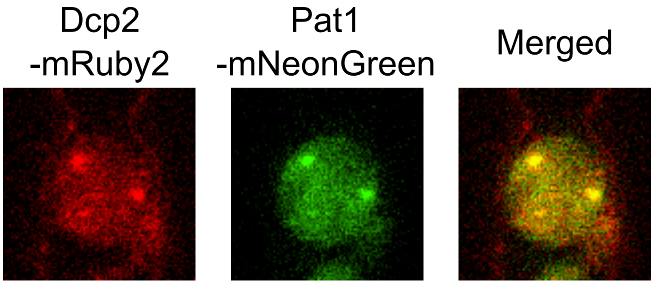


**Figure S2. A representative image of colocalized Pat1 and Dcp2 in a wild type cell measured in a microfluidics device.** The age of the cell shown here was the 12th division.

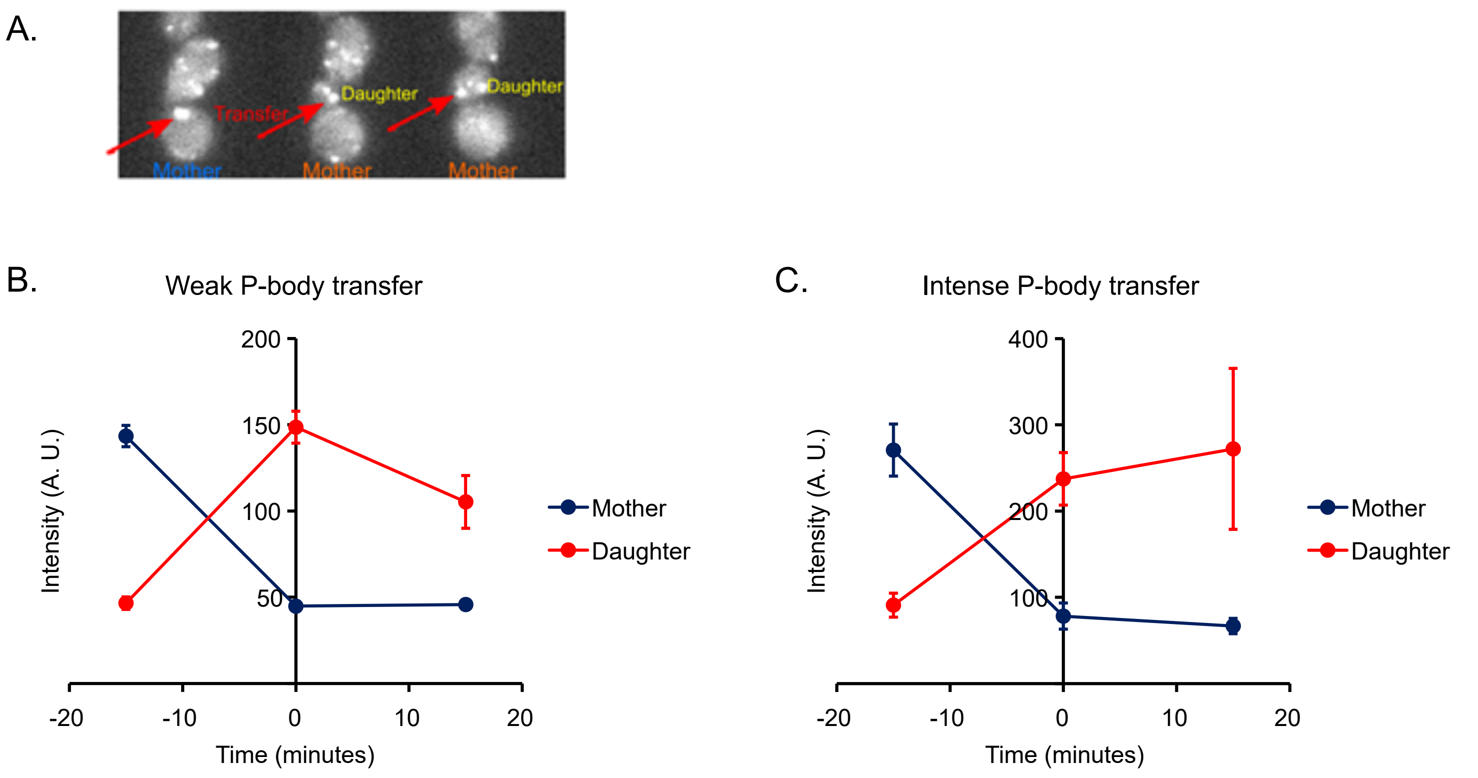


**Figure S3. Intensity profile of P-body transfer. A.** A representative image of P-body transfer from a mother to a daughter cell. Mother cells indicated in blue and orange are the ones before and after P-body transfer, respectively. Red arrow indicates a transferred P-body. **B-C.** Changes in maximum intensities of the cytoplasm when the weak and the intense P-bodies were transferred from mother to daughter cells, respectively (mean ± S. E.). P-body transfer happened at time 0.
